# High-density genetic linkage mapping in Sitka spruce advances the integration of genomic resources in conifers

**DOI:** 10.1101/2023.08.21.554184

**Authors:** Hayley Tumas, Joana J. Ilska, Sebastien Girardi, Jerome Laroche, Stuart A’Hara, Brian Boyle, Mateja Janes, Paul McLean, Gustavo Lopez, Steve J. Lee, Joan Cottrell, Gregor Gorjanc, Jean Bousquet, John A. Woolliams, John J. MacKay

## Abstract

In species with large and complex genomes such as conifers, dense linkage maps are a useful for supporting genome assembly and laying the genomic groundwork at the structural, populational and functional levels. However, most of the 600+ extant conifer species still lack extensive genotyping resources, which hampers the development of high-density linkage maps. In this study, we developed a linkage map relying on 21,570 SNP makers in Sitka spruce (*Picea sitchensis* [Bong.] Carr.), a long-lived conifer from western North America that is widely planted for productive forestry in the British Isles. We used a single-step mapping approach to efficiently combine RAD-Seq and genotyping array SNP data for 528 individuals from two full-sib families. As expected for spruce taxa, the saturated map contained 12 linkages groups with a total length of 2,142 cM. The positioning of 5,414 unique gene coding sequences allowed us to compare our map with that of other Pinaceae species, which provided evidence for high levels of synteny and gene order conservation in this family. We then developed an integrated map for *P. sitchensis* and *P. glauca* based on 27,052 makers and 11,609 gene sequences. Altogether, these two linkage maps, the accompanying catalog of 286,159 SNPs and the genotyping chip developed herein opens new perspectives for a variety of fundamental and more applied research objectives, such as for the improvement of spruce genome assemblies, or for marker-assisted sustainable management of genetic resources in Sitka spruce and related species.

## Introduction

Recombination frequency analysis was developed over a century ago to order genetic markers (Sturevant 1913), leading to the development of genetic linkage maps and ultimately the linking of phenotypic traits to chromosomal regions. Genetic linkage mapping (e.g., Gyapay et al. 1994), along with high-throughput DNA sequencing was instrumental in producing the first human genome sequence assembly (IHGSC 2001). In plants, linkage maps allowed for positioning gene coding regions and anchoring sequence scaffolds obtained through whole genome sequencing (WGS) in a variety of species including poplar (Tuskan et al. 2006), potato (Xu et al. <u>2011</u>), *Eucalyptus* (Myburg et al. 2014), ryegrass (Velmurugan et al. <u>2016</u>), soybean (Song et al. <u>2016),</u> or spruces (Gagalova et al. 2022). Linkage maps are useful for laying the genomic groundwork in species with genomes that are difficult to assemble due to size or complexity, such as barley (5.1 Gb) and wheat (16 Gb) (Mascher *et al,* 2013; Chapman *et al*. 2015), which both have large hexaploid genomes and abundant repetitive sequences. For this reason, the development of the first saturated linkage maps in conifers (e.g., Devey et al. 1994; Pelgas et al. 2005), which have very large genomes (18-34 Gb) and extensive repetitive regions (MacKay et al. 2012; De La Torre et al. 2014), predates by two decades the report of first genome assemblies (Birol et al. 2013; Nystedt et al. 2013; Zimin et al. 2014; Warren et al. 2015). Despite the rapid development of sequencing technologies, genetic linkage maps remain an essential genomic resource for species with such large genomes and highly fragmented genome assemblies (De La Torre et al. 2014). The importance of a wide number of conifer species in breeding programs and productive forestry across the globe (Mullin et al. 2011) has encouraged the development of genetic linkage maps and other genomic resources to support fundamental research and diverse applications (Bousquet et al. 2021).

One of the main findings emerging from comparative genome mapping studies in conifers has been the detection of high levels of intergeneric macro-synteny and macro-collinearity among Pinaceae taxa (Pelgas et al. 2006; Ritland et al. 2011; Pavy et al. 2012; Westbrook et al. 2015). The low incidence of large chromosomal rearrangements, despite the ancient divergence within the group, has enabled the development of consensus maps across species. For example, the high structural conservation in *Pinus taeda* L. and *Pinus elliottii* Engelm. enabled the development of a consensus genetic map with 3856 markers (Westbrook et al. 2015), and similarly across *Picea glauca* (Moench) Voss and *Picea mariana* (Mill.) B.S.P. (Pavy et al. 2008, 2012). Likewise, highly conserved gene coding sequences among Pinaceae taxa has allowed to transfer efficiently exome capture sequencing probes across species, for instance, probes originally developed in *P. glauca* (Sena et al. 2014) were successfully used for large-scale SNP discovery in gene coding regions of *P. mariana* (Pavy et al. 2016) and *Picea abies* (L.) H. Karst (Azaiez et al. 2018).

However, most of the 600+ extant conifer species still lack linkage maps or have maps with a low marker density, limiting their usefulness in molecular breeding applications or other genomic analyses. Nonetheless, the recent advent of high throughput genotyping techniques has allowed to use DNA markers covering thousands of genetic loci, and to develop high-density linkage maps in a number of conifer and plant species. As for most forest trees, conifers have high levels of genetic diversity and heterozygosity (Hamrick and Godt, 1990), which has facilitated the large-scale discovery of single nucleotide polymorphisms (SNPs) by expression tag sequencing (Dantec et al. 2004; Pavy et al. 2006), targeted resequencing by using exome capture (Neves et al, 2014; Pavy et al. 2016, Azaiez et al. 2018), and genotyping-by-sequencing (GBS) (e.g., Gamal El-Dien et al. 2015). As a result, extensive genomic resources have been developed but only for a few commercially-relevant Pinaceae taxa, such as *P. taeda* L. (Neves et al. 2014), *Pinus pinaster* Aiton (Plomion et al. 2015; de Miguel et al. 2015), *Pinus flexilis* (E. James) Rydberg (Liu et al. 2019), *P. glauca* (Pavy et al. 2013, 2017), *P. mariana* (Pavy et al. 2016) and *P. abies* (Bernhardsson et al. 2019). Several genotyping methods have been used in conifers, from custom chips (Moriguchi et al, 2012; Pavy et al. 2008, 2013, 2016; Plomion et al. 2015) to targeted sequencing (Neves et al. 2014; Bernhardsson et al. 2019) and reduced representation whole-genome sequencing (e.g., restriction site associated DNA sequencing (El-Dien et al. 2015)).

The reported high levels of genome synteny and collinearity among the Pinaceae provide an opportunity to accelerate the development of genomic resources in ecologically and economically relevant species such as Sitka spruce (*Picea sitchensis* [Bong.] Carr.), for which a large database of mRNA sequences (Ralph et al. 2008) and a draft genome assembly (Gagalova et al. 2022) are available, but still lacks a large-scale genotyping resource or a high-density linkage map. *P. sitchensis* is a long-lived conifer found mostly in the coastal areas of western North America, and that is widely planted for forestry in the British Isles (Lee et al. 2013). Linkage mapping and genomic selection are of great interest to improve our understanding of the genetic basis of quantitative traits appropriate for tree breeding (Lee et al. 2013; Fuentes-Utrilla et al. 2017), and genetic diversity management to maintain or increase resilience to damaging pests in *P. sitchensi*s in the context of exotic forestry and climate change (Tumas et al, 2021). Here, we aimed to develop genomic markers that could be used in conjunction with a comparative genomic approach to produce genetic linkage. Our specific objectives were as follows: 1) Use probes developed in *P. glauca* (Stival Sena et al. 2014) to perform exome capture and SNP discovery in Sitka spruce; 2) Develop a large-scale SNP array for genotyping natural and mapping Sitka populations; 3) Develop high-density linkage maps from full-sib (FS) families by using data from the SNP array and previous restriction site associated DNA sequencing (RAD-Seq) data; 4) Compare the resulting *P. sitchensis* linkage map to maps from those available for other conifers; and 5) Develop an integrated *Picea* genetic map with *P. glauca*.

## Material and Methods

### Study Population, Sampling and DNA Extraction

All plant materials in this study were from two distinct *P. sitchensis* full-sib genetic field trials (Trial 1 and Trial 2) established in the United Kingdom. Information on the trials is presented in Figure 1, along with details on samples used for 1) SNP discovery (orange), 2) SNP Chip validation (green), or 3) linkage map development (blue), and which samples in Trial 1 had additional RAD-seq genotyping data (Fuentes-Utrilla *et al*. 2017) used in linkage map development (Figure 2). Briefly, Trial 1, consisting of three full-sib families replicated across three sites, was used for SNP discovery and linkage map development while Trial 2 comprised 50 full-sib families across two sites and was used for SNP discovery and validation in this study and to develop genomic prediction in a separate study (Ilska et al *in revision*). Samples from two full-sib families in Trial 1 (Family 1: SS1773 x SS3159, Family2: SS493 x SS1463) were used in linkage map development and were all collected from a single site in Llandovery, UK (Fuentes-Utrilla *et al*. 2017), for genotyping using either the SNP Chip, RAD-Seq, or both methods (Figure 1).

**Figure 1.**
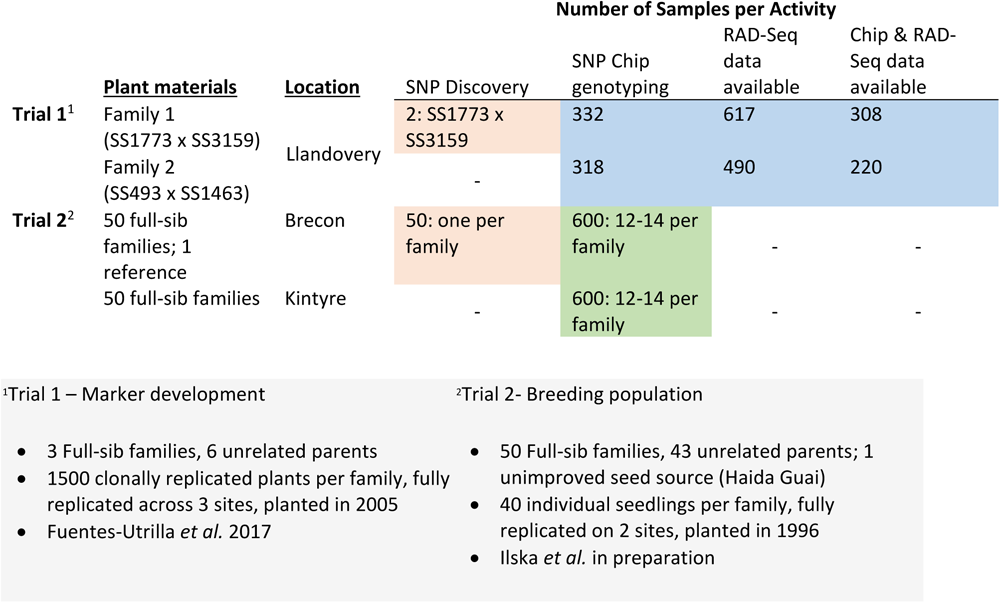
Details of the full-sib genetic trials (in grey) and the samples taken from these which were used in SNP discovery (orange) for the SNP Chip and SNP Chip validation (green), the subset of samples used for SNP Chip and RAD-Seq genotyping to develop the linkage map (blue).

**Figure 2.**
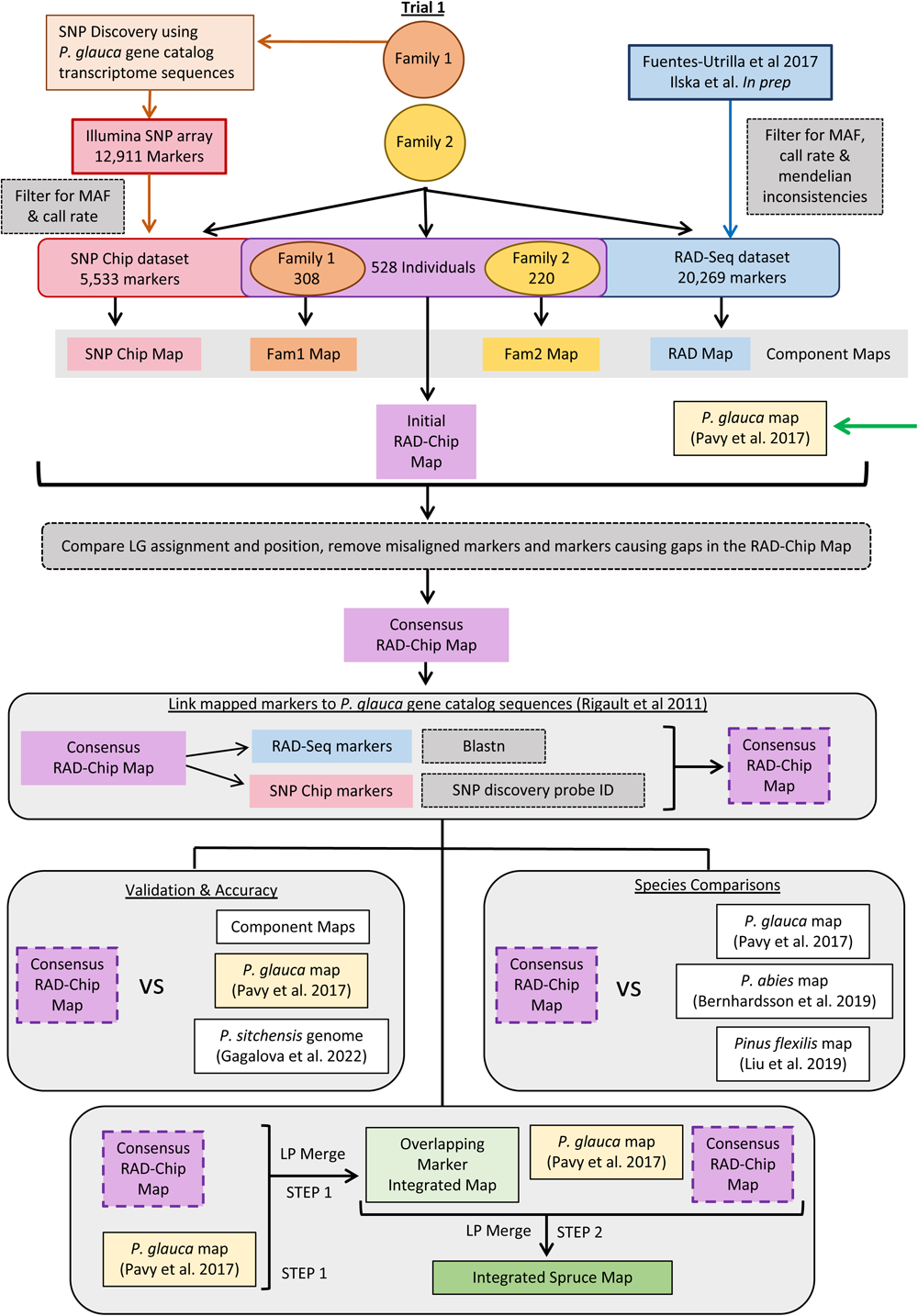
Schematic the diagram of the Sitka spruce map development, validation, species comparison and map integration steps. Several Sitka spruce component maps were initially developed using each family and marker type; however, the final RAD-Chip map was developed as a single step combining both families and both the RAD-Seq and the Infinium Chip data (Top). Comparative genomic analysis approaches were then used to assess the accuracy of the map, conduct to study synteny across species and then develop an integrated spruce map (Bottom).

The sampling for SNP discovery and SNP chip genotyping was carried out in July-August 2017, whereas the RAD-Seq sampling was completed as described previously in Fuentes-Utrilla et al. (2017). All samples were comprised of foliage from healthy annual growth collected by removing 1-3 shoot tips of approximately 5-10 cm in length from healthy branches, subsequently placing them intact in sealed, labeled plastic bags, and storing them in a cool box for less than 48 hours. After transporting them to the laboratory the needles were removed from the rachis and stored at -20°C until used for DNA isolation.

DNA was isolated for SNP discovery from parents of the linkage mapping families in Trial 1 and one randomly selected individual for each of the 50 full-sib families in Trial 2 by Forest Research. Needles (100 mg) were finely chopped, placed in 2 ml Eppendorf tubes with two 3 mm stainless-steel ball bearings, and ground to a fine powder in a Retsch mixer-mill (Retsch, Haan, Germany). DNA was isolated from powder using a Qiagen DNAeasy Plant mini-kit (QIAGEN, Hilden, Germany) with the following modifications. Lysis buffer AP1 volume was increased from 400 to 600µl and incubation time was increased from 10 to 20 minutes. Neutralization buffer (P3) volume was increased from 130 to 200µl and a constant volume of 800µL of AW1 wash buffer was added to each sample. During the elution step, eluted product was re-applied to the column, incubated for five minutes, and then spun down to elute the final product. DNA concentration was measured using a Qubit fluorometer (original model, ThermoFisher Scientific, Massachusetts, USA). DNA for the SNP Chip was isolated from 50 mg of needles by the Austrian Institute of Technology (AIT, Vienna Austria). DNA isolation for RAD-Seq genotyping was as described in Fuentes-Utrilla et al. (2017).

### Exome Capture Sequencing

Samples from two parents of one family in Trial 1 (SS1773 x SS3159) and a pool of samples comprised of one individual of each of the 50 families in Trial 2 (Figure 1) were used as libraries for SNP discovery to develop the SNP chip (Figure 2). The Trial 2 pool was assembled by merging untagged extracted DNAs from each individual in equimolar concentration. One large insert (avg. 650 bp) NebNext Ultra II library (New England Biolabs, Ipswich, MA) was generated for each of the two Trial 1 parents (SS1773 and x SS3159) and for the pool of Trial 2 samples, following the manufacturer’s instructions.

Oligonucleotide probes used herein to capture *P. sitchensis* gene homologs were originally designed from *P. glauca* transcriptome sequences (Rigault et al. 2011) and were previously used successfully under an exome capture framework on *P. glauca* (Stival Sena et al. 2014), *P. mariana* (Pavy et al. 2016), and *P. abies* (Azaiez et al. 2018). Multiple probes (0.5M) ranging from 50 to 105 nucleotides in length were designed for each of 23,684 transcripts of the white spruce GCAT catalog, with each base being covered by two probes on average. Two micrograms of libraries (100 ng from each parent, 900 ng from the breeding population) were used in a liquid-phase capture (SeqCap EZ developer, IRN 6089042357, OID35086, Roche Nimblegen). The captured material was amplified and sequenced on an Illumina HiSeq 4000 PE100 at the Centre d’Expertises et de Services Génome Québec (Montréal, QC, Canada). Illumina HiSeq 4000 sequencing generated two ∼ 100-bp paired-end sequences per captured insert, which yielded over 403M raw sequences for the three libraries (Supp Table 1).

**Table 1.** Library sequencing and read mapping summary statistics (Nb=Numbers of)

| **Library** | **Nb. raw reads** | **Nb. reads post quality control** | **Nb. reads post mapping** | **Supplementary\*** | **Nb. mapped reads (%)** | **Nb. unmapped reads (%)** |
| --- | --- | --- | --- | --- | --- | --- |
| Trial 1 Parent SS1773 | 100,001,026 | 99,811,140 | 99,973,099 | 161,959 | 24,576,905 (25) | 75,396,194 (75) |
| Trial 1 Parent SS3159 | 64,519,124 | 64,092,588 | 64,196,564 | 103,976 | 15,883,403 (25) | 48,313,161 (75) |
| Trial 2 families | 238,330,524 | 237,123,494 | 237,538,801 | 415,307 | 62,800,361 (26) | 174,738,440 (74) |
| Total | 402,850,674 | 401,027,222 | 401,708,464 | 681,242 | 103,260,669 (26) | 298,447,795 (74) |

\* For reads that aligned in a chimeric fashion, one segment was designated as primary, and the remainder as supplementary.

**Table 2.** Summary of SNP Chip genotyping results across trials and SNP discovery populations. Informative SNPs were polymorphic after filtering for call rate and MAF (see methods); uninformative SNPs gave a low call rate, were monomorphic or polymorphic with a low MAF. (Nb=Numbers of)

| **SNP discovery population** | **Nb on Chip** | **Trial 1 - informative** | **Trial 1 - Uninformative** | **Trial 2 - informative** | **Trial 2 - Uninformative** |
| --- | --- | --- | --- | --- | --- |
| Recycled | 1554 | 1407 (95%) | 147 (5%) | 1466 (94%) | 88 (6%) |
| Mapping Parents | 5303 | 1828 (34%) | 3475 (66%) | 2375 (45%) | 2928 (55%) |
| 50 F-S families | 6054 | 2298 (38%) | 3756 (62%) | 3105 (51%) | 2949 (49%) |
| ALL | 12911 | 5533 (43%) | 7378 (57%) | 6946 (54%) | 5965 (46%) |

### Read Libraries Processing, Reference-guided Alignment, and SNP Detection

For the reads obtained for each library, Illumina adapter sequences were removed using the software Cutadapt 2.7 (Martin 2011), and sequencing quality was checked before and after adapter removal with the software FastQC Version 0.11.8 (Andrews 2010). After this step, 100M,64M, and ∼237M sequences were obtained for the two Trial 1 libraries (SS1773 and SS3159) and Trial 2 library, respectively (Supp Table 1). Reads were then mapped to the most complete version of the white spruce (*P. glauca*) catalog of expressed genes GCAT3.3 (Rigault et al. 2011), which contains 27,720 gene cluster sequences. *P. sitchensis* and *P. glauca* are closely-related taxa which can hybridize (Hamilton et al. 2013), therefore, this strategy allowed to maximize gene representation and facilitate subsequent SNP selection for the design of a genotyping array (see ‘Genotyping assay’ section below for further details). Each library was aligned to the reference genome using the BWA-MEM algorithm (Li et al. 2010) and was converted to BAM format with SAMtools (Li et al. 2011). Around 25% of the sequences mapped to the reference genome, representing a total of over 100M mapped sequences (Supp Table 1). Next, variant calling was performed with the software Platypus v0.8.1 (Rimmer et al. 2014). The minimum number of supporting reads required to consider a variant was set to 25, and all remaining criteria were Platypus default parameters. Variant calling with Platypus resulted in the identification of 286,159 SNPs distributed across 23,480 GCAT gene clusters.

### SNP Chip Assay and Genotyping

The SNPs discovered here were used along with 1,554 SNPs already identified in *P. sitchensis* from a previous genotyping study using *P. glauca* Infinium genotyping arrays (Pavy et al. 2013) to develop a new Infinium iSelect array (Illumina, San Diego, CA) for genomic analyses. All newly discovered SNPs retained for building the array met the following general criteria: 1) were strictly biallelic SNPs; 2) included only one SNP per gene and were type II SNPs (one bead per SNP) whenever possible, in order to maximize the number of genes represented on the array; 3) carried no SNP or indel within 50 bp in their 5’ or 3’ flanking regions (Illumina probe design requirement); and 4) had Illumina functionality scores ≥ 0.4. More specifically, SNPs observed in at least one mapping parent library (SS1773 or SS3159) were retained when they met the following criteria: depth ≥ 25; Minor Allele Frequency (MAF) ≥ 0.25 in Trial 1 parent libraries, and MAF ≥ 0.05 in the Trial 2 library. SNPs observed in the Trial 2 library only were also selected when their MAF exceeded 0.15, and their depth exceeded 50 reads. For this last subset of SNPs, when more than one SNP was available for a given gene, the SNP with MAF around 0.25 was retained so as to filter out possible paralogs expected to yield MAFs close to 0.5.

Following chip manufacture by Illumina, genotypes were obtained both for the mapping families from Trial 1 (analysed in this report) and for the full-sib of Trial 2 (analysed in a separate study in preparation) at the Centre d’expertise et de services Génome Québec (Montréal, QC, Canada, group of Daniel Vincent). Genotype calling was performed using the GenomeStudio v2.0.5 software (Illumina), and genotype clusters were visually examined and manually curated when necessary to reject monomorphic and failed polymorphisms. Excel files output from GenomeStudio were formatted for PLINK v1.90b4 (Purcell et al. 2007) using R v4.0.2 (R Core Team, 2022). Data for the Trial 1 individuals were filtered separately in PLINK to retain only those SNPs with a minimum call rate of 80% and a minor allele frequency (MAF) greater than 0.2, and to exclude individuals with a call rate below 85%, then data were reformatted into variant calling files (VCF). The data for the Trial 2 individuals were also filtered in PLINK –mind 0.05 and MAF greater than 0.05.

### Combining Datasets

Data from the Infinium iSelect SNP array (SNP Chip dataset) were combined with a dataset from a previous study (Fuentes-Utrilla et al. 2017; Ilska et al. in revision) that used RAD-seq to genotype a similar set of samples from the same full-sib families in Trial 1 (RAD-seq dataset) (Figure 1). Following filtering for individual (60%) and SNP (80%) call rate, MAF (0.15) and mendelian inconsistencies, the RAD-seq dataset contains 15,452 and 17,915 genotyped loci across 617 and 490 offspring and parents for family 1 and 2, respectively. Samples across the two families were combined in the RAD-Seq dataset and a single SNP per locus was chosen based on call rate for a total of 27,967 SNPs across 1,111 individuals. The SNP Chip and RAD-Seq datasets were joined by overlap in sampled individuals with 308 and 220 individuals, including parents, overlapping in family 1 and 2, respectively for a total of 528 individuals in the combined dataset (Figure2). These overlapping individuals were extracted from VCFs containing each complete SNP Chip and RAD-Seq dataset using VCFtools v0.1.16 (Danecek et al. 2011) and then resulting VCFs were merged using ‘concat’ in BCFtools v1.8 (Danecek et al. 2021) to combine datasets for mapping.

### Constructing Linkage Maps

Linkage maps were constructed using Lep-MAP3 v0.2 (Rastas 2017) with java v8.45.14. Lep-MAP3 allows family data to be combined and used simultaneously for construction of linkage maps. In total, five maps were constructed, one consensus map that is the main resource of this study and four component maps developed from different subsets of the data that were used for method validation (Figure 2). The consensus map (RAD-Chip Map) used both families 1 and 2 and the combined SNP Chip and RAD-Seq marker dataset. The four component maps were constructed separately, two using both families but each of the marker datasets separately (SNP Chip and RAD maps) and two using the combined marker dataset but only samples from each family separately (Fam1 and Fam2 maps, corresponding to Family 1 and Family 2 in Figure 1). Data for each of these five maps was input into Lep-MAP3 using ‘ParentCall2’ allowing the removal of noninformative markers (removeNonInforamtive=1) and then filtered using ‘Filtering2’ using the default data tolerance for segregation distortion of 0.01. Markers were assigned to linkage groups (chromosomes) using ‘SeparateChromosomes2’, testing a range of minima for the logarithm of the odds (LOD) score between groups of markers (lodLimit) between 15 and 95 and with a minimum of 100 markers set as the requirement to form a group. When developing the two family component maps (Fam1 and Fam2), Fam2 had a linkage group length distribution in centimorgans (cM) more comparable to that found in *P. glauca* (Pavy et al. 2017) while Fam1 had a much longer first linkage group exceeding 247 cM. For this reason, data for family 2 was used to inform marker grouping for the other three maps that combined families (RAD-Chip, SNP Chip, and RAD maps) using the ‘families’ function within ‘SeparateChromosomes2’. Markers that were not assigned to a linkage group were then added to these generated linkage groups using ‘JoinSingles2All’ by testing a range of lodLimits from 2 to 50 and using a lodDifference of 10. The best lodLimit was selected for each step by determining which value assigned the most markers to 12 linkage groups, the known number of chromosomes in *P. sitchensis* (Supplemental Table 1). Markers were ordered on linkage groups and relative position in cM was determined using ‘OrderMarkers2’ with the Kosambi mapping function (useKosambi=1) and averaging marker position over sex (sexAveraged=1). This ordering step was iterated five times for each chromosome and the order with the highest likelihood was selected as the final map for each dataset or family.

The consensus RAD-Chip map was further developed by removing problematic markers, which were identified by examining gaps at the end of linkage groups and by checking for inconsistencies in linkage group assignment and order of markers or genes in the RAD-Chip map compared to the four other component maps (SNP Chip, RAD, Fam1, Fam2) as well as the *P. glauca* map (Pavy et al. 2017). Gaps were identified visually by plotting the RAD-Chip map in ggplot2 (Wickham 2016) in R v4.03 (R Core Team 2022). Differences in linkage group assignments were determined by merging resulting maps and aggregating by linkage group in R v4.03. Marker order was compared between maps using linear models with the ‘lm’ function in R v4.03, based on the idea that, when plotted against one another, marker positions within a linkage group should have a linear relationship when maps have similar marker order. Using only markers that grouped the same across the two maps, the position of the marker in the component or *P. glauca* map was regressed against the position in the RAD-Chip map for each linkage group. Cook’s distance (Cook 1977) was used to identify any markers that had a substantially different position in the two maps, using a threshold of 4/n, where n is the number of markers in the comparison. Lep-MAP3 was rerun on the RAD-Chip dataset as described above using only those markers that mapped previously, excluding markers that caused gaps at the end of linkage groups, were assigned to different linkage groups in two or more map comparisons, or markers that surpassed the Cook’s distance threshold in any comparison.

### Map Validation and Accuracy

Marker linkage group assignment and order in the consensus RAD-Chip map was validated against the four component maps (SNP Chip, RAD, Fam1, Fam2) by calculating the proportion of markers assigned to the same linkage group and the correlation in marker order using Kendall’s τ in R v4.03. Accuracy of the RAD-Chip map linkage group assignment and marker ordering was verified using the *P. glauca* gene catalog GCAT3.3 (Rigault et al. 2011) and the *P. sitchensis* genome sequence (Gagalova et al. 2022, Genbank assembly no. GCA_010110895.2). Map assignment and ordering were considered accurate if markers located on the same *P. glauca* gene or *P. sitchensis* genome scaffold were assigned to the same linkage group and located at the same position or within a window of 10 cM to one another. All SNP Chip SNPs have a corresponding *P. glauca* gene from using the *P. glauca* transcriptome sequences to inform exome capture in the SNP discovery. RAD-Seq SNPs were matched to the *P. glauca* catalog using Blastn (Altschul et al. 1990, Camacho et al. 2008), calling reciprocal best hits with a minimum of 95% identity and a maximum E-value of 1×10^-11^. Sequences of *P. glauca* catalog genes with matches in the RAD-Chip map were used in Blastn to find matches in the *P. sitchensis* genome using reciprocal best hits with a minimum of 95% identity and a maximum E-value of 1×10^-100^. Linkage group and position were compared among any markers that were located on the same sequence or scaffold.

### Comparisons to Other Species

Synteny across other species within the *Pinaceae* family was examined using a consensus map from Norway spruce (*Picea abies* (L.) Karst.) (Bernhardsson et al. 2019) and maps of *P. glauca* (Pavy et al. 2017) and limber pine (*Pinus flexilis* James) (Liu et al. 2019)(Figure 2). The set of sequences from the *P. glauca* gene catalog (Rigault et al. 2011) matching to either SNP Chip markers or RAD-Seq markers on the RAD-Chip map, determined either during the SNP discovery for the SNP Chip markers or with Blastn as described above for the RAD Seq markers, were used as the basis of comparison to the other species. Mapped markers in the *P. glauca* map all sit within a sequence in the *P. glauca* gene catalog (Rigault et al. 2011, Pavy et al. 2013, Pavy et al. 2017), allowing for direct comparison. For comparison to *P. abies* and *P. flexilis*, Blastn was used to find orthologous genes between *P. sitchensis* and the mapped genes in each species. Markers mapped in *P. abies* were identified using sequence capture based on the *P. abies* genome assembly v1.0 (Nystedt et al. 2013) available on ConGenIE (www.congenie.org) and markers mapped in *P. flexilis* were identified using sequence capture based on a *P. flexilis* transcriptome provided by the Liu et al. (2019) authors. Mapped sequences were extracted from these files for synteny analysis. Orthologous marker pairs were identified as reciprocal best hits with a maximum E-value of 1×10^-100^ and a minimum of 95% and 90% identity when comparing to *P. abies* genome scaffolds and *P. flexilis* transcriptome sequences, respectively. In the particular case when orthologous *P. abies* SNPs were located on the same genome scaffold, but assigned to different linkage group according to the *P. abies* map, both SNPs were included in the synteny analyses and accounted for statistically. Synteny was evaluated both visually and statistically by calculating the proportion of orthologous genes that were assigned to the same linkage group and estimating the correlation in the marker order using Kendall’s τ.

### Constructing an Integrated Spruce Map

Using the *P. glauca* gene catalog (Rigault et al. 2011) annotation associated with the *P. sitchensis* SNP Chip dataset and the markers on the *P. glauca* map along with the additional matches in the mapped RAD-Seq markers as described above, *P. sitchensis* and *P. glauca* (Pavy et al. 2017) maps were integrated with the aim of developing a spruce map that included more genes than either of the species maps alone (Figure 2). To simplify integration of gene placement across species and prevent inconsistencies among markers, a single marker per *P. glauca* gene catalog sequences was used. The *P. glauca* map only includes one marker per catalog gene, so first, a single marker per gene catalog sequence was selected from the markers with a match to the catalog on the RAD-Chip map, preferentially selecting SNP Chip markers. Second, markers causing major discrepancies between the two maps were removed, i.e., markers that either were not assigned to the same linkage group or markers that had surpassed the threshold for order misalignment using Cook’s distance as described above. Filtered maps were then combined using ‘LPMerge’ (Endelman and Plomion 2014) in R v4.03 in two steps.

In the first step, only markers found on both maps were used to create an integrated map, giving the maps weights equivalent to sample size and testing a maximum interval size between bins from 1 to 10 across each linkage group. The second step generated the final integrated species map by combining the filtered individual species maps with all markers with the resulting merged map from the first step, giving the filtered RAD-Chip and *P. glauca* maps and the merged map weights of one, two, and three respectively to reflect sample size and confidence. In both steps the best consensus map was selected across the interval size bins by both comparing the lowest root mean-squared error (RMSE) for each linkage group and by comparing the integrated species map linkage group length to the average length of the RAD-Chip and *P. glauca* map linkage groups. In the integrated species map from step 2 containing all possible markers, large gaps were created at the ends of linkage groups. The markers creating these gaps were manually removed in the final integrated species map. Synteny with the component species maps and this final integrated species map was assessed by calculating the percentage of markers assigned to the same linkage group, the correlation in marker order using Kendall’s τ, and visually through graphs in R.

## Results

### SNP Chip Genotyping

The final SNP Chip array contained 12,911 markers across 12,893 unique sequences from the *P. glauca* gene catalog (see methods for identification and selection of SNPs). Of those markers, 1,554 were previously shown to be polymorphic in *P. sitchensis* according to an array designed for *P. glauca* (Pavy et al. 2013) and 4,604 had been previously mapped in *P. glauca* (Pavy et al. 2017). Following filtering for call rate and MAF (see methods for details), 5,533 markers were successfully called as informative across the two linkage mapping families, among which 2,572 markers were from the previously mapped in *P. glauca*. Similarly, 6,946 markers were successfully called as informative in a total of 1262 individuals from Trial 2 trees.

### Linkage Maps and Map Integrity

In the SNP Chip dataset, three and eight offspring were removed from families 1 and 2 respectively, due to low call rate and 5,533 markers across the two families passed filtering for call rate and MAF in PLINK. Following further filtering in Lep-MAP3 for segregation distortion and uninformative markers, 615 samples (including of parents) and 5,194 markers in total were retained to develop the SNP Chip component map. Additional filtering in Lep-MAP3 reduced the RAD-Seq dataset from 27,967 to 19,529 markers across 1,111 samples in both families for use in the RAD component map. The combined SNP Chip and RAD-Seq datasets contained 25,802 markers, 5,533 from the SNP Chip dataset and 20,269 from the RAD-Seq dataset, across the 528 individuals genotyped with both methods. Following additional filtering in Lep-MAP3, 24,702 markers were used to construct the RAD-Chip consensus map, 5,194 from the SNP Chip dataset and 19,508 from the RAD-Seq dataset. The component family maps used 14,499 and 15,955 markers across 308 and 220 samples for Fam 1 and 2, respectively, following filtering in Lep-MAP3.

As expected, all maps placed markers across 12 linkage groups. The component SNP Chip, RAD, Fam1, and Fam2 maps placed 5,064, 15,041, 12,685, and 14,831 markers respectively, with an average total map length of 2,412.3 cM that ranged from 2,148.2 for the RAD map to 2,927.6 for the SNP Chip map. The initial RAD-Chip consensus map placed 22,505 markers with a total map length of 2,367.3 cM. A total of 934 markers were excluded from these 22,505 markers for causing gaps or consistent differences in assignment or order across comparisons to the component maps and the *P. glauca* map. The finalized RAD-Chip map that excluded these problematic markers mapped 21,570 markers for a total map length of 2,141.6 cM and an average distance of 0.1 cM between markers.

In comparison to the four component maps, 99.90-99.99% of markers were assigned to the same linkage group in the finalized RAD-Chip map with the least agreement occurring with the Fam1 map (Figure S1). Concordance in marker order ranged from 0.95-0.99 with the lowest correlation occurring with the Chip (Figure S1B). When linking mapped markers to the *P. glauca* catalog, 4,590 genes were mapped from the SNP Chip dataset and 1,094 were mapped from the RAD-Seq dataset, which resulted in a total of 5,414 unique genes positioned on the final map (270 overlapping genes between both datasets). Of these genes, 326 were linked to two or more markers for a total of 670 markers located on genes carrying at least one other marker. Across these 670 markers, 92% were assigned to the same linkage group as the co-occurring SNP with an average distance of 3 cM (0-157.2 cM) between co-occurring markers. When linking mapped markers to the *P. sitchensis* genome using *P. glauca* catalog sequences, 2,161 unique scaffolds were represented on the final map with 32 scaffolds matching more than one marker arising from the co-occurrence of 70 markers. Only two scaffolds had markers that were not assigned to the same linkage group with 94% of co-occurring markers assigned to the same linkage group and an average distance of 0.8 cM (0-2.7 cM) between co-occurring markers.

### Synteny Across Species

Synteny between the *P. sitchensis* consensus RAD-Chip map and *P. glauca* map (Pavy et al. 2017) was based on 2,581 catalog genes. These genes corresponded to 2,778 marker pairs on both maps with 94% of them being assigned to the same linkage group and with an average concordance in marker order of 0.98 (0.96-0.98) across linkage groups (Figure 3B). A total of 3,234 marker pairs were used to compare the *P. sitchensis* map to the *P. abies* consensus map (Bernhardsson et al. 2019), corresponding to 1,935 *P. glauca* catalog genes and 1,873 *P. abies* genome scaffolds (Figure 3A,B). Across marker pairs, 88% were assigned to the same linkage group with an average correlation in marker order of 0.96 (0.91-0.98). Synteny with the *P. flexilis* (Limber pine) map (Liu et al. 2019) was based on 1,514 marker pairs with 1,397 unique *P. flexilis* markers on 1,397 unique *P. glauca* catalog genes with 85% of markers assigned to the same linkage group and an average correlation in marker order of 0.93 (0.88-0.97) (Figure 3C). In all three comparisons, markers that did not align were distributed evenly across the 12 linkage groups (Figure 3) and there were no indications of inversions or translocations in marker order within linkage groups (Figure 3).

**Figure 3.**
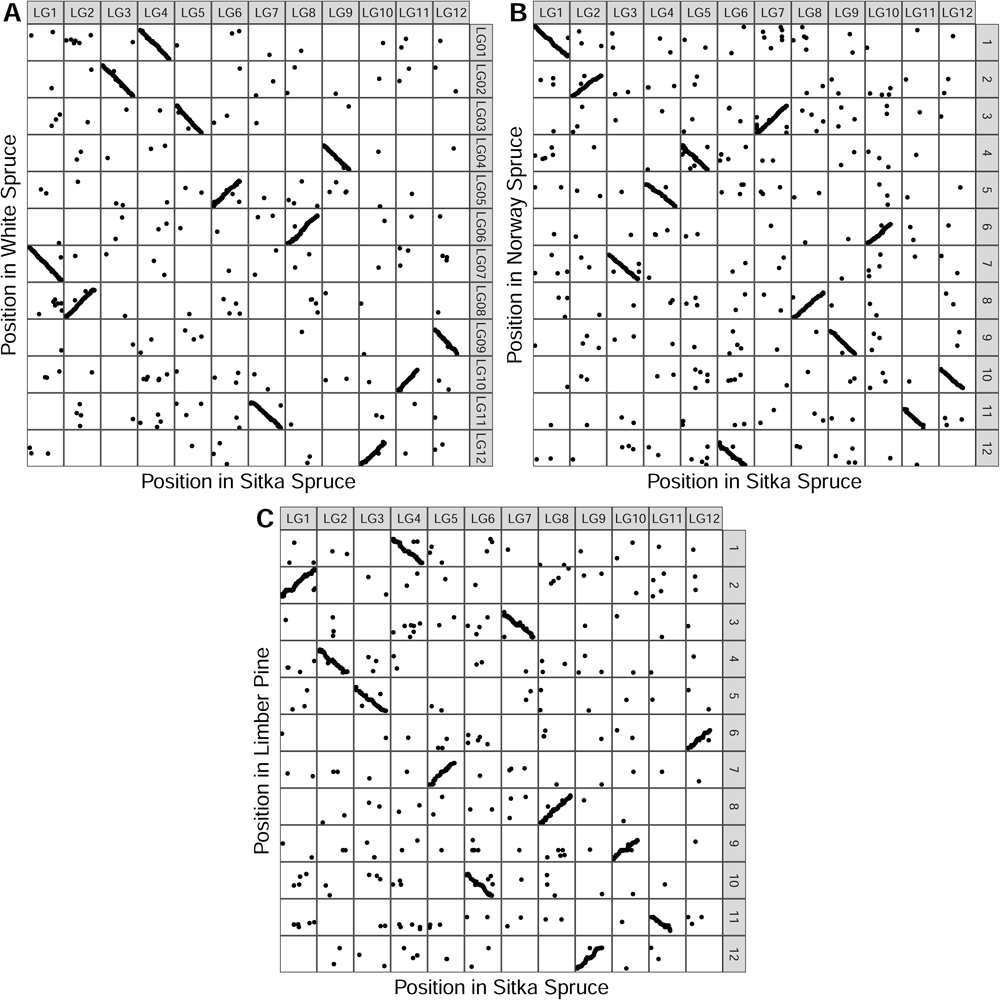
Comparison of marker assignment and order on chromosomes (linkage groups) between Sitka spruce (x-axis) and linkage maps in three other species from previous studies (y-axis): **A** white spruce (Pavy et al. 2017), **B** Norway spruce (Bernhardsson et al. 2019), and **C** limber pine (Liu et al. 2019). Note that the linkage group (LG) labels on the y-axis are taken from the originally published map in each species.

### An Integrated Spruce Map

A total of 2,327 markers that were present in both the *P. glauca* and *P. sitchensis* maps were used to produce a merged map of overlapping markers. After selecting a single marker per *P. glauca* catalog gene from the *P. sitchensis* map and filtering for discrepancies between the *P. sitchensis* and *P. glauca* maps, 20,983 and 8,539 markers were selected to integrate the *P. sitchensis* and *P. glauca* maps, respectively, including 2,327 overlapping markers. After removing gaps at the terminal ends of linkage groups, the final integrated map contained 27,052 markers with 11,331 *P. glauca* catalog genes for a total map length of 1,860.2 cM and an average distance between markers of 0.07 cM. The integrated map placed an additional 6,195 *P. glauca* catalog genes compared to the *P. sitchensis* map and an additional 18,519 markers, including 2,798 additional *P. glauca* catalog genes, compared to the *P. glauca* map (Figure 4A). All markers were assigned to the same linkage groups in both maps and concordance in marker order averaged 0.96 and 0.99 across linkage groups for the *P. sitchensis* and *P. glauca* maps respectively (Figure 4B,C).

**Figure 4.**
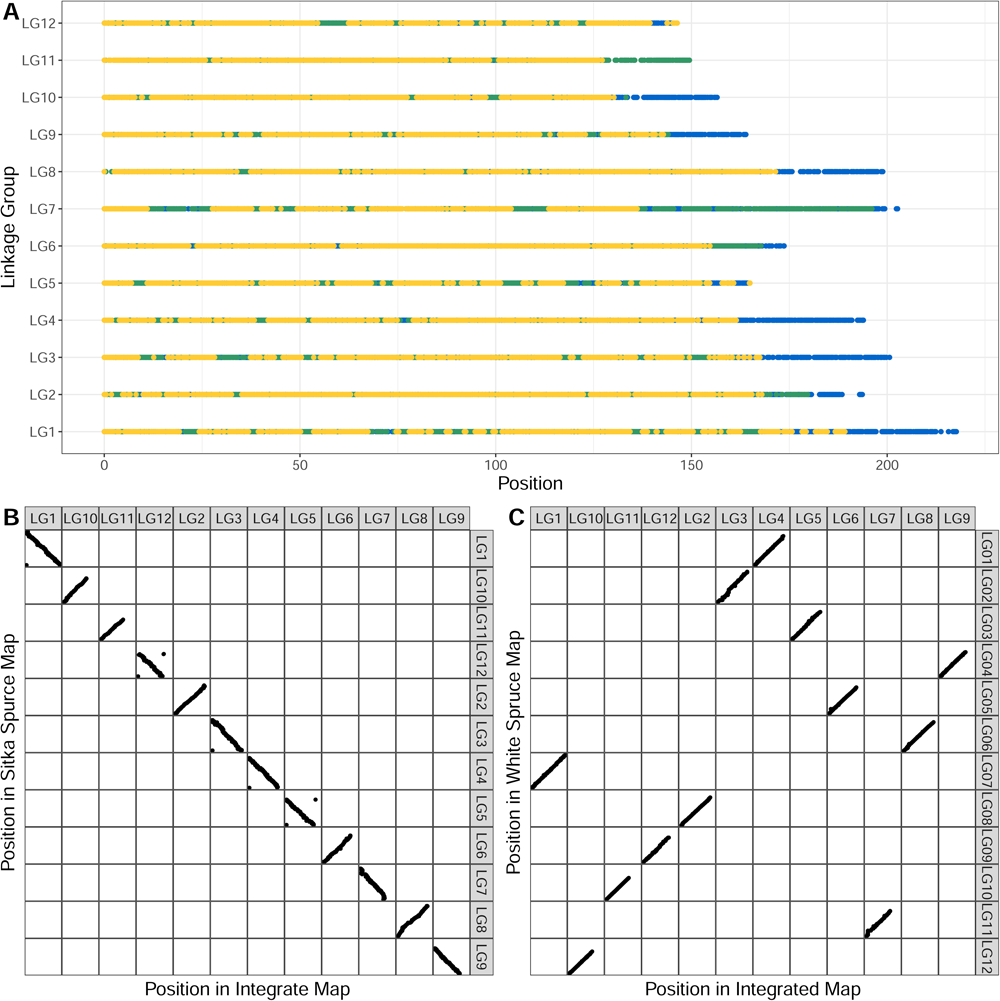
A representation of the integrated map of Sitka and white spruce and comparison to individual species maps. **A** Chromosome or linkage group (LG) is on the y-axis with marker position on the x-axis. Points represent markers that are color-coded for markers found in both species maps that were used to create an initial map of only overlapping markers (yellow) and markers found only in Sitka spruce (green) or white spruce (blue) that were added in a second integration step to make the consensus integrated map. **B,C** Comparison of marker assignment and order on linkage groups between the Sitka spruce (**B**) and white spruce (**C**) maps (x-axis) to the integrated map (y-axis).

## Discussion

The size and complexity of conifer genomes has limited the assembly of high-quality whole genome sequences, as indicated by the high degree of fragmentation in early whole genome assemblies (e.g. Nystedt et al., 2013; Birol et al., 2013; Zimin et al. 2014; De La Torre et al. 2014; Warren et al. 2015). Not surprisingly, conifer genome assemblies are still only available for species of high economic or ecological significance, and population-level genome resequencing is generally lacking. The importance and utility of linkage maps to assist assignments of large scaffolds to linkage groups was recently illustrated in a comparative genomic study focusing on *Picea* species, in conjunction with the use of long-read sequencing methods (Gagalova et al. 2022). Our *P. sitchensis* map is comparable in terms of the number of mapped genes (5414) and superior based on the density of markers (21,570) to other recently expanded conifer genetic maps that integrated markers obtained by next-generation sequencing, such as those made for *P. glauca* (Gagalova et al. 2022; 14,727 expressed genes), *P. abies* (Bernhardsson et al. 2019 – 21,506 markers containing 17,079 gene models), *P. flexilis* (Liu et al. 2019 – 9,612 gene models, Maritime pine (*P. pinaste*r) (Chancerel et al. 2013 -∼1100 markers), and loblolly pine (*P. taeda*) (Westbrook et al. 2015 - 3856 markers across 3305 genome scaffolds). Here, we discuss how our approach has simplified the development and integration of maps, and highlight how the resulting resource can expand our understanding of conifer genomes and support genetic resource management.

### Map Development and Integration

In this study, by using two types of genotypic data and maps from two species we were able to maximise the number of markers we mapped and, gather and integrate a large amount of genomic information. This was made possible by combining RAD-Seq and SNP-array genotypic data for the same individuals of two unrelated Sitka spruce full-sib families. This allowed us to merge and order both marker types together, while also combining family data on the front-end during map development, a feature unique to LepMap-3 (Rastas 2017). Earlier studies have relied on combining multiple marker types in forest trees but most often on a smaller scale. For example, AFLPs, ESTPs, SSRs and gene-based SNPs were mapped together in *P. glauca* (831 markers, 348 genes) and in *P. mariana* (835 markers, 328 genes) (Pavy et al. 2008). Alternatively, several distinct maps produced with different marker types were reconstructed using gene-based SNPs, based on the same principles as applied here (Westbrook et al. 2015). In contrast, we obtained and used large SNP datasets from both exome sequencing and RAD-Seq, and by analyzing two independent full-sib families. This allowed us to produce individual component maps to verify for map coherence across genetic backgrounds before producing a consensus map with all data combined (Figure 2). Recent high-density genetic maps have used a simpler approach based on a single marker type (Neves et al. 2014; Plomion et al. 2015; de Miguel et al. 2015; Pavy et al. 2017; Liu et al. 2019; Bernhardsson et al. 2019); however, our approach allowed us to draw inferences efficiently across marker types in a single step without requiring a map integration step.

By using an exome capture probe set designed and validated in *P. glauca* (Stival et al. 2014) for SNP discovery in *P. sitchensis*, we explicitly aimed to obtain genotyping data in overlapping genomic sequences to enable direct comparisons across multiple conifer species. While this approach has been successfully used previously for SNP discovery across species, it had not yet been used to create an integrated map. The 4,590 array-SNP markers and 1,094 RAD-Seq markers were located in or matched a *P. glauca* transcriptome sequence, allowing us to compare our *P. sitchensis* consensus map robustly with the *P. glauca* map (Pavy et al. 2017). The gene-based markers also allowed comparison with linkage maps in *P. abies* (Bernhardsson et al. 2019) and *P. flexilis* (Liu et al. 2019). This comparison indicated highest levels of synteny in Picea-Picea comparisons, with levels ranging as expected owing to respective pairwise phylogenetic distance, i.e., slightly lower synteny in the Picea-Pinus comparison. This is also the first study to integrate high density linkage maps from two different conifer species, creating a resource that is more informative for each individual species. The integrated map provides information on conserved gene locations across species and provides a foundation for further development and integration with other species towards a more complete and comprehensive conifer genomic resource.

### Evolutionary Insights and Resource for Breeding and Conservation

We developed a high-density linkage map with 21,570 makers in *P. sitchensis* and an integrated map with 27,052 markers for *P. sitchensis* and *P. glauca*, both of which should facilitate further improvement of conifer genome sequence assemblies and contiguity. There is a high level of macro-synteny and macro-collinearity among species in the Pinaceae (e.g. Pavy et al. 2012; Westbrook et al. 2015), and this apparent lack of chromosomal rearrangement enables genomic integration across species such as the creation of consensus genetic maps, as seen for *P. taeda* and *P. elliottii* with 3856 shared markers (Westbrook et al. 2015). Many of the current conifer genome assemblies are still highly fragmented (e.g., Nystedt et al. 2013; Zimin et al. 2014; Gagalova et al. 2022) and contain many partial gene sequences (e.g., Warren et al. 2015), which leaves large gaps in our ability to conduct comparative genomics and evolutionary studies.

Recently, the integration of an expanded high-density linkage map from *P. glauca* and shotgun genome assemblies was reported in *P. glauca*, *P. engelmanii*, *P. sitchensis,* and a natural hybrid of the three species (interior spruce) (Gagalova et al. 2022). Up to 32% of genome scaffolds could be anchored to linkage groups and further assembled into super-scaffolds representative of chromosomes, in addition to validating those areas of the genome assembly (Gagalova et al. 2022). Up to 65% of genomic scaffolds could be recently anchored on the *P. glauca* genetic map following improvement of genome assemblies using longer reads (R Warren and I Birol, University of British Columbia, personal communication). Therefore, the integrated linkage map generated herein will further improve this rate and inform genome assemblies more exhaustively, particularly for *P. sitchensis,* and facilitate cross-species comparisons among *Picea* spp. This will result in an improved structural characterization of conifer genomes including micro-rearrangements and the organization of genes in tandem arrays or functional operons (Pavy et al. 2017). This augmented spruce consensus linkage map has the potential to shed new insights on early lineage divergence and their genomic footprints in the conifers, such as between the Pinaceae and Cupressaceae (Moriguchi et al. 2012, de Miguel et al. 2015) and Taxaceae families, with the recent release of genome assemblies for *Sequoiadendron giganteum* (Lindl.) J. Buchh (Scott et al. 2020), Sequoia *sempervirens* Endl. (Neale et al. 2021), and *Taxus chinensis* (Pilger) Rehd. (Xiong et al. 2021).

We have discovered 286,159 SNPs distributed across 23,480 *P. sitchensis* gene clusters after exome capture and sequencing, with probes developed on *P. glauca* (Stival et al. 2014), and previously validated in *P. mariana* (Pavy et al. 2016) and *P. abies* (Azaiez et al. 2018). The highly conserved nature of gene coding sequences across these congeneric species made it possible to successfully transfer probes between taxa and suggests that the set of probes used in this study should work across the whole *Picea* genus. By targeting common genes across species, our SNP discovery approach allowed us to develop a genotyping array which selectively included genes that were both unmapped and previously mapped in *P. glauca* (Pavy et al. 2017). This approach aimed to facilitate direct cross-species comparisons by using previously mapped genes as anchoring points, and filling gaps in previous maps by positioning unmapped genes. This strategy has resulted in an integrated map including 27,052 markers, which will offer an opportunity to increase our understanding of gene family evolution in conifers and plants more broadly. For example, in conifers, stress related gene families have been reportedly more diverse than in Angiosperms (Rigault et al. 2011; Warren et al. 2015; De la Torre Cuba et al. 2020; Gagalova et al. 2022), and a high level of sequence novelty was found across conifer species in dehydrin (Stival et al. 2018), NLR (Van Ghelder et al. 2019), and R2R3-MYB (Bedon et al. 2010) gene families, among others. In the conifer *P. flexilis*, linkage mapping showed that disease resistance NLR genes were highly clustered on a few linkage groups (Liu et al. 2019). Therefore, new linkage maps and integrated genomic resources (as reported here or in Gagalova et al. 2022) should help to further our understanding of the evolution of this large gene family.

From an applied perspective, the resources developed herein will also support genetic resource management in *P. sitchensis*, the dominant productive forestry species in the British Isles. These resources include a genotyping array of 12,911 SNPs, the acquisition of large-scale genotypic data for two mapping families and a breeding population (Figure 1), as well as high-density linkage maps. Genomic selection is poised to accelerate and transform forest tree breeding, although its implementation in breeding programs targeting both gymnosperm and angiosperm trees remains challenging (Grattapaglia et al. 2022). In conifers such as *Pinus spp.* and *Picea spp.*, genomic selection gave genetic prediction abilities approaching those achieved with pedigree-based selection for growth, wood properties and insect resistance (Beaulieu et al. 2014; Lenz et al. 2020; Bousquet et al. 2021; Calleja-Rodriguez et al., 2020; Isik et al. 2022). However, the high cost associated with the acquisition of large-scale genotypic and phenotypic data still represents a barrier for routine use of genomic prediction in tree breeding programs (Klápště et al. 2022). In a companion study to this report, Ilska et al. (in revision) have used the newly-developed *P. sitchensis* linkage map to impute missing RAD-Seq genotypes in mapping families 1 and 2, and in one unrelated family, which resulted in improved call rates by up to 10%, and a significant reduction of genotyping costs by allowing the use of lower-cost genotyping methods, which performance in genomic selection schemes is generally negatively affected by lower call rates and genome coverage. The genotypes obtained in the current *P. sitchensis* breeding population are currently analysed in a distinct genomic selection study of growth and wood traits (Ilska et al. in revision) and in another related study aiming at developing a low-cost DNA-fingerprinting assay (MacKay et al. in preparation), following Godbout et al. (2017).

The linkage maps presented here will also facilitate mapping quantitative trait loci (QTL) for traits related to adaptation (Pelgas et al. 2011; Pavy et al. 2017; Laoué et al. 2021) and pest resistance (Lind et al. 2014), among others, and comparative studies of genomic architecture to better understand the divergent or convergent nature of the evolution across spruces and other conifers. It would also allow to identify new candidate genes for further investigations at the functional level or for diagnostic marker development. Finally, evolutionary studies using linkage maps to gain insights into the structure of large gene families involved in disease resistance such as nucleotide-binding and leucine-rich repeat (NLR) genes (Van Ghelder et al. 2019; Liu et al. 2019), or dehydrin genes for drought response (Stival Sena et al. 2018) may result in potent diagnostic tools and benefit forest practitioners involved in the management and the conservation of genetic resources in natural and breeding populations, especially in the context of accelerated climate change.

## Conclusion

In this study, we developed a highly densified genetic linkage map in *P. sitchensis* by efficiently combining different marker types and by targeting gene coding regions to facilitate comparative genomic analyses and integration across species (Figure 2). Together, the newly identified SNP markers and new genetic linkage maps will help improving genome assemblies, expanding our understanding of conifer genome evolution, and supporting *P. sitchensis* genomic resource development and genetic resource management. Gymnosperms have been reported to have less diverse and less dynamic genomes compared to those of flowering plants (Leitch and Leitch. 2012) but interestingly, they are genetically diverse and have a large proportion of their rapidly evolving genes related to stimuli and stress response (Gagalova et al. 2022), many of which belong to highly diversified gene families (e.g., Bedon et al. 2010; Stival Sena et al. 2018; Van Ghelder et al. 2019). Conifers are also characterized by high levels of intraspecific phenotypic variability in defensive compounds (e.g., Mageroy et al. 2015; Parent et al. 2020; Tumas et al. 2021). The resources reported here will aid the understanding, conservation and sustainable use of this wealth of adaptive potential to support the resilience of natural and breeding populations in the face of climate change.

## Data Availability

Genotype tables for each map (i.e., RAD, Chip, and RAD-Chip maps), pedigree files, code to convert files to LepMap3 format, final map files for the three maps and the integrated map, and SNP array information are available at Dryad (*pending* DOI). Code described to develop linkage maps, compare composite maps and species maps, and develop the integrated map are publicly available at: https://github.com/HayleyTumas/SitkaLinkageMap.

## Acknowledgments

We thank David Gil-Moreno, Andy Price, Robert Sykes (Forest Research), Glyndwr Jones (University of Oxford) for technical assistance with the foliage sampling. Natural Resources Wales provided in kind support with the provision of materials and forest operations. Rajesh Joshi provided help with LepMap scripts. This work was funded by the Biotechnology and Biological Sciences Research Council (Grant #: BB/PZ020488/1) as part of the Sitka Spruced project. Support is also acknowledged from the Spruce-Up project supported by Genome Canada, Genome Quebec, and Genome British Columbia (Grant # : 234FOR).

## Literature cited

1. Altschul SF, Gish W, Miller W, Myers EW, Lipman DJ. 1990. Basic local alignment search tool. J. Mol. Biol. 215:403–410. 10.1016/S0022-2836(05)80360-2

2. Andrews S. 2010. FastQC: A Quality Control Tool for High Throughput Sequence Data. https://github.com/s-andrews/FastQC

3. Azaiez A, Pavy N, Gérardi S, Laroche J, Boyle B, Gagnon F, Mottet M-J, Beaulieu J, Bousquet J. 2018. A catalog of annotated high-confidence SNPs from exome capture and sequencing reveals highly polymorphic genes in Norway spruce (Picea abies). BMC Genomics 19: 942. 10.1186/s12864-018-5247-z

4. Beaulieu J, Doerksen TK, MacKay J, Rainville A, Bousquet J.2014. Genomic selection accuracies within and between environments and small breeding groups in white spruce. BMC Genomics 15:1048. 10.1186/1471-2164-15-1048

5. Bedon F, Bomal C, Caron S, Levasseur C, Boyle B, Mansfield SD, Schmidt A, Gershenzon J, Grima-Pettenati J, Séguin A, MacKay J.2010. Subgroup 4 R2R3-MYBs in conifer trees: gene family expansion and contribution to the isoprenoid-oriented response. Journal of Experimental Botany 61: 3847–3864. 10.1093/jxb/erq196

6. Bernhardsson C, Vidalis A, Wang X, Scofield DG, Schiffthaler B, Baison J, Street NR, García-Gil MR, Ingvarsson PK. 2019. An Ultra-Dense Haploid Genetic Map for Evaluating the Highly Fragmented Genome Assembly of Norway Spruce (Picea abies), G3 Genes|Genomes|Genetics 9(5): 1623–1632. 10.1534/g3.118.200840

7. Birol IA, Raymond SD, Jackman S, Pleasance R, Coope GA, Taylor MM, Saint Yuen CI, Keeling D, Brand BP, Vandervalk H, Kirk P, Pandoh RA, Moore Y, Zhao AJ, Mungall B, Jaquish, B, Yanchuk A, Ritland C, Boyle B, Bousquet J, Ritland K, MacKay J, Bohlmann J, Jones SJM. 2013. Assembling the 20 Gb white spruce (Picea glauca) genome from whole-genome shotgun sequencing data. Bioinformatics 29:1492–1497. 10.1093/bioinformatics/btt178

8. Calleja-Rodriguez A, Pan J, Funda T, Chen Z, Baison J, Isik F, Abrahamsson S, Wu HX. 2020. Evaluation of the efficiency of genomic versus pedigree predictions for growth and wood quality traits in Scots pine. BMC Genomics. 21:796. 10.1186/s12864-020-07188-4

9. Camacho C, Coulouris G, Avagyan V, Ma N, Papadopoulos J, Bealer K, Madden TL. 2008. BLAST+: architecture and applications. BMC Bioinformatics 10:421. 10.1186/1471-2105-10-421

10. Chancerel E, Lamy JB, Lesur I, Noirot C, Klopp C, Ehrenmann F, Boury C, Le Provost G, Label P, Lalanne C, Léger V, Salin F, Gion J-M, Plomion C. 2013. High density linkage mapping in a pine tree reveals a genomic region associated with inbreeding depression and provides clues to the extent and distribution of meiotic recombination. BMC Biology 11:50. 10.1186/1741-7007-11-50

11. Chapman JA, Mascher M, Buluç A, Barry K, Georganas E, Session A, Strnadova V, Jenkins J, Sehgal S, Oliker L, Schmutz J, Yelick KA, Scholz U, Waugh R, Poland JA, Muehlbauer GJ, Stein N, Rokhsar DS. 2015. A whole-genome shotgun approach for assembling and anchoring the hexaploid bread wheat genome. Genome Biol. 16:26. 10.1186/s13059-015-0582-8

12. Danecek P, Auton A, Abecasis G, Albers CA, Banks E, DePristo MA, Handsaker RE, Lunter G, Marth GT, Sherry ST, McVean G, Durbin R. 2011. The variant call format and VCFtools. Bioinformatics 27(15):2156–8. 10.1093/bioinformatics/btr330

13. Danecek P, Bonfield JK, Liddle J, Marshall J, Ohan V, Pollard MO, Whitwham A, Keane T, McCarthy SA, Davies RM, Li H. 2021. Twelve years of SAMtools and BCFtools. Gigascience 10(2):giab008. 10.1093/gigascience/giab008

14. de Miguel M, Bartholomé J, Ehrenmann F, Murat F, Moriguchi Y, Uchiyama K, Ueno S, Tsumura Y, Lagraulet H, de Maria N, Cabezas JA, Cervera MT, Gion JM, Salse J, Plomion C. 2015. Evidence of intense chromosomal shuffling during conifer evolution. Genome Biol Evol 7(10):2799–2809. 10.1093/gbe/evv185

15. De la Torre AR, Piot A, Liu B, Wilhite B, Weiss M, Porth I. 2020. Functional and morphological evolution in gymnosperms: a portrait of implicated gene families. Evolutionary Applications, 1(13):210–227. 10.1111/eva.12839

16. Devey ME, Fiddler TA, Liu BH, Knapp SJ, Neale DB. 1994. An RFLP linkage map for loblolly pine based on a three-generation outbred pedigree. Theor Appl Genet 88(3-4):273–8. 10.1007/BF00223631. PMID: 24186005

17. Fuentes-Utrilla P, Goswami C, Cottrell JE, Pong-Wong R, A’Hara SW, Woolliams JA. 2017. QTL analysis and genomic selection using RADseq derived markers in Sitka spruce: the potential utility of within family data. Tree Genetics & Genomes 13:33. 10.1007/s11295-017-1118-z

18. Gagalova K, Warren RL, Coombe L, Wong J, Nip KM, Saint Yuen MM, Whitehill J, Celedon JM, Ritland C, Taylor GA, Cheng D, Plettner P, Hammond SA, Mohamadi H, Zhao Y, Moore RA, AMungall AJ, Boyle B, Laroche J, Cottrell J, Mackay J, Lamothe M, Gérardi S, Isabel N, Pavy N, Jones SJM, Bohlmann J, Bousquet J, Birol I. 2022. Spruce giga-genomes: structurally similar yet distinctive with differentially expanding gene families and rapidly evolving genes. The Plant Journal 111(5):1469–1485. 10.1111/tpj.15889

19. Gamal El-Dien O, Ratcliffe B, Klápště J, Chen C, Porth I, El-Kassaby YA. 2015. Prediction accuracies for growth and wood attributes of interior spruce in space using genotyping-by-sequencing. BMC Genomics 16:370. 10.1186/s12864-015-1597-y

20. Godbout J, Tremblay L, Levasseur C, Lavigne P, Rainville A, Mackay J, Bousquet J, Isabel N. 2017. Development of a traceability system based on a SNP array for large-scale production of high-value white spruce (Picea glauca). Frontiers in Plant Science 8:1264. 10.3389/fpls.2017.01264

21. Grattapaglia D. 2022. Twelve Years into Genomic Selection in Forest Trees: Climbing the Slope of Enlightenment of Marker Assisted Tree Breeding. Forests 13(10):1554. 10.3390/f13101554

22. Gyapay G, Morissette J, Vignal A, Dib C, Fizames C, Millasseau P, Mard S, Bernardi G, Lathrop M, Weissenbach J. 1994. The 1993–94 Généthon human genetic linkage map. Nat Genet 7:246–339. 10.1038/ng0694supp-246

23. Hamilton JA, Lexer C, Aitken SN. 2013. Genomic and phenotypic architecture of a spruce hybrid zone (Picea sitchensis × P. glauca). Molecular Ecology 22:827–841. 10.1111/mec.12007.

24. Hamrick JL, Godt MJ. 1990. Allozyme diversity in plant species. In Plant population Genetics, Breeding, and Genetic Resources, Eds. Brown AHD, Clegg MT, Kahler AL, Weir BS. Sinauer Associates, Sunderland, MA. pp. 43–63.

25. Ilska JJ, Tolhurst DJ, Tumas H, Maclean P, Cottrell J, Lee SJ, Mackay J, Woolliams JA (2023) Additive and non-additive genetic variance in juvenile Sitka spruce (Picea sitchensis, Bong. Carr), In revision, Tree Genes and Genomes.

26. International Human Genome Sequencing Consortium (IHGSC).2001. Initial sequencing and analysis of the human genome. Nature 409:860–921. 10.1038/35057062

27. Isik F. 2022. Genomic Prediction of Complex Traits in Perennial Plants: A Case for Forest Trees. In Complex Trait Prediction: Methods and Protocols, Eds. Ahmadi N, Bartholomé J, Humana: New York, NY, USA. pp. 493–520. 10.1007/978-1-0716-2205-6_18.

28. Klápště J, Ismael A, Paget M, Graham NJ, Stovold GT, Dungey HS, Slavov GT. 2022. Genomics-Enabled Management of Genetic Resources in Radiata Pine. Forests 13:282. 10.3390/f13020282

29. Lee S, Thompson D, Hansen JK. 2013 Sitka spruce (Picea sitchensis (Bong.) Carr). In Forest tree breeding in Europe: current state-of-the-art and perspectives, Managing Forest Ecosystems, vol25 Eds. Pâques LE. Springer, Dordrecht, NL, pp. 177–227. 10.1007/978-94-007-6146-9_4.

30. Leitch AR, Leitch IJ. 2012. Ecological and genetic factors linked to contrasting genome dynamics in seed plants. New Phytologist 194:629–646. 10.1111/j.1469-8137.2012.04105.x

31. Lenz PRN, Nadeau S, Mottet MJ, Perron M, Isabel N, Beaulieu J, Bousquet J. 2020. Multi-trait genomic selection for weevil resistance, growth, and wood quality in Norway spruce. Evol. Appl. 13:76–94. 10.1111/eva.12823

32. Li Z, Baniaga AE, Sessa EB, Scascitelli M, Graham SW, Rieseberg LH, Barker MS. 2015. Early genome duplications in conifers and other seed plants. Sci Adv. 1(10):e1501084. 10.1126/sciadv.1501084

33. Li H, Durbin R. 2010. Fast and accurate long-read alignment with Burrows-Wheeler Transform. Bioinformatics 26: 589–595. 10.1093/bioinformatics/btp698

34. Li H. 2011. A statistical framework for SNP calling, mutation discovery, association mapping and population genetical parameter estimation from sequencing data. Bioinformatics 27: 2987–93. 10.1093/bioinformatics/btr509

35. Liu J-J, Schoettle AW, Sniezko RA, Yao F, Zamany A, Williams H, Rancourt B. 2019. Limber pine (Pinus flexilis James) genetic map constructed by exome-seq provides insight into the evolution of disease resistance and a genomic resource for genomics-based breeding. Plant J, 98: 745–758. 10.1093/10.1111/tpj.14270

36. Mackay J, Dean JFD, Plomion C, Peterson DG, Cánovas FM, Pavy N, Ingvarsson PK, Savolainen O, Guevara MA, Fluch S, Vinceti B, Abarca D, Díaz-Sala C, Cervera M-T. 2012. Towards decoding the conifer giga-genome. Plant Molecular Biology 80: 555-569. 10.1007/s11103-012-9961-7

37. Mageroy MH, Parent GJ, Germanos G, Giguère I, Delvas N, Maaroufi H, Bauce É, Bohlmann J, Mackay J. 2015. Expression of the beta-glucosidase gene Pgβglu-1 underpins natural resistance of white spruce against spruce budworm. Plant Journal 81(1):68–80. 10.1111/tpj.12699

38. Martin M. 2011. Cutadapt removes adapter sequences from high-throughput sequencing reads. EMBnet.journal 17: 10–12. 10.14806/ej.17.1.200

39. Mascher M, Muehlbauer GJ, Rokhsar DS, Chapman J, Schmutz J, Barry K, Muñoz-Amatriaín M, Close TJ, Wise RP, Schulman AH, Himmelbach A, Mayer KF, Scholz U, Poland JA, Stein N, Waugh R. 2013. Anchoring and ordering NGS contig assemblies by population sequencing (POPSEQ). Plant J. 76:718–727. 10.1111/tpj.12319

40. Moriguchi Y, Ujino-Ihara T, Uchiyama K, Futamura N, Saito M, Ueno S, Matsumoto A, Tani N, Taira H, Shinohara K, Tsumura Y (2012) The construction of a high-density linkage map for identifying SNP markers that are tightly linked to a nuclear-recessive major gene for male sterility in Cryptomeria japonica D. Don.BMC Genomics. Mar 16; 13: 95.

41. Myburg AA, Grattapaglia D, Tuskan GA, Hellsten U, Hayes RD, Grimwood J, Jenkins J, Lindquist E, Tice H, Bauer D, Goodstein DM, Dubchak I, Poliakov A, Mizrachi E, Kullan AR, Hussey SG, Pinard D, van der Merwe K, Singh P, van Jaarsveld I, Silva-Junior OB, Togawa RC, Pappas MR, Faria DA, Sansaloni CP, Petroli CD, Yang X, Ranjan P, Tschaplinski TJ, Ye CY, Li T, Sterck L, Vanneste K, Murat F, Soler M, Clemente HS, Saidi N, Cassan-Wang H, Dunand C, Hefer CA, Bornberg-Bauer E, Kersting AR, Vining K, Amarasinghe V, Ranik M, Naithani S, Elser J, Boyd AE, Liston A, Spatafora JW, Dharmwardhana P, Raja R, Sullivan C, Romanel E, Alves-Ferreira M, Külheim C, Foley W, Carocha V, Paiva J, Kudrna D, Brommonschenkel SH, Pasquali G, Byrne M, Rigault P, Tibbits J, Spokevicius A, Jones RC, Steane DA, Vaillancourt RE, Potts BM, Joubert F, Barry K, Pappas GJ, Strauss SH, Jaiswal P, Grima-Pettenati J, Salse J, Van de Peer Y, Rokhsar DS, Schmutz J. 2014. The genome of Eucalyptus grandis. Nature 510:356–362. 10.1038/nature13308

42. Neale DB, Zimin AV, Zaman S, Scott AD, Shrestha B, Workman RE, Puiu D, Allen BJ, Moore ZJ, Sekhwal MK, De La Torre AR, McGuire PE, Burns E, Timp W, Wegrzyn JL, Salzberg SL. 2022. Assembled and annotated 26.5 Gbp coast redwood genome: a resource for estimating evolutionary adaptive potential and investigating hexaploid origin. G3 (Bethesda). 12(1): jkab380. 10.1093/g3journal/jkab380

43. Neves LG, Davis JM, Barbazuk WB, Kirst M. 2014. A high-density gene map of loblolly pine (Pinus taeda L.) based on exome sequence capture genotyping. G3 (Bethesda) 4(1): 29–37. 10.1534/g3.113.008714

44. Niu S, Li J, Bo W, Yang W, Zuccolo A, Giacomello S, Chen X, Han F, Yang J, Song Y, Nie Y, Zhou B, Wang P, Zuo Q, Zhang H, Ma J, Wang J, Wang L, Zhu Q, Zhao H, Liu Z, Zhang X, Liu T, Pei S, Li Z, Hu Y, Yang Y, Li W, Zan Y, Zhou L, Lin J, Yuan T, Li W, Li Y, Wei H, Wu HX. 2022. The Chinese pine genome and methylome unveil key features of conifer evolution. Cell 185(1):204–217. 10.1016/j.cell.2021.12.006

45. Nystedt N, Street NR, Wetterbom A, Zuccolo A, Lin Y-C, Scofield DG, Vezzi F, Alexeyenko A, Giacomello S, Delhomme N, Vicedomini R, Sahlin K, Sherwood E, Elfstand M, Gramzow L, Holmberg,K, Keech O, Klasson L, Koriabine M, Kucukoglu M, Käller M, Luthman J, Lyshol, F, Niittylä T, Olson Å, Ritland C, Rosselló JA, Sena J, Svensson T, Talavera-López T, Theißen G, Tuominen H, Vanneste K, Wu Z-Q, Zhang B, Zerbe P, Arvestad L, Bhalerao R, Bohlmann J, Bousquet J, Garcia Gil J, Hvidsten TR, de Jong P, MacKay J, Morgante M, Ritland K, Sundberg B, Lee Thompson S, Van de Peer Y, Andersson B, Nilsson O, Ingvarsson PK, Lundeberg J, Jansson S. 2013. Sequencing the 20 Gbp Norway spruce genome sheds light on conifer genome evolution. Nature 497: 579–84. 10.1038/nature12211

46. Parent GJ, Méndez-Espinoza C, Giguère I, Melissa H. Mageroy MH, Martin Charest M, Bauce E, Bohlmann, J, MacKay JJ. 2020. Hydroxyacetophenone defenses in white spruce against spruce budworm. Evolutionary Applications 13: 62– 75. 10.1111/eva.12885

47. Pavy N, Lamothe M, Pelgas B, Gagnon F, Birol I, Bohlmann J, Mackay J, Isabel N, Bousquet J. 2017. A high-resolution reference genetic map positioning 8.8K genes for the conifer white spruce: Structural genomics implications and correspondence with physical distance. The Plant Journal 90:189–203. 10.1111/tpj.13478

48. Pavy N, Pelgas B, Laroche J, Rigault P, Isabel N, Bousquet J. 2012. A spruce gene map infers ancient plant genome reshuffling and subsequent slow evolution in the gymnosperm lineage leading to extant conifers. BMC Biology 10: 84. 10.1186/1741-7007-10-84

49. Pavy N, Gagnon F, Deschênes A, Boyle B, Beaulieu J, Bousquet J. 2016. Development of highly reliable in silico SNP resource and genotyping assay from exome capture and sequencing: an example from black spruce (Picea mariana). Molecular Ecology Resources 16 588–598. 10.1111/1755-0998.12468

50. Pavy, N, Gagnon F, Rigault P, Blais S, Deschênes A, Boyle B, Pelgas B, Deslauriers M, Clément S, Lavigne P, Lamothe M, Cooke JEK, Jaramillo-Correa JP, Beaulieu J, Isabel N, Mackay J, Bousquet J. 2013. Development of high-density SNP genotyping arrays for white spruce (Picea glauca) and transferability to subtropical and nordic congeners. Molecular Ecology Resources 13: 324–336. 10.1111/1755-0998.12062

51. Plomion, C, Bartholomé J, Lesur I, Boury C, Rodríguez-Quilón I, Lagraulet H, Ehrenmann F, Bouffier L, Gion JM, Grivet D, de Miguel M, de María N, Cervera MT, Bagnoli F, Isik F, Vendramin GG, González-Martínez SC. 2016. High-density SNP assay development for genetic analysis in maritime pine (Pinus pinaster). Molecular Ecology Resources, 16: 574–587. 10.1111/1755-0998.12464

52. R Core Team. 2022. R: A language and environment for statistical computing. R Foundation for Statistical Computing, Vienna, Austria. URL https://www.R-project.org/

53. Rastas P. 2017. Lep-MAP3: robust linkage mapping even for low-coverage whole genome sequencing data. Bioinformatics 33(23):3726–3732. 10.1093/bioinformatics/btx494

54. Rigault P, Boyle B, Lepage P, Cooke JEK, Bousquet J, MacKay J. 2011. A white spruce gene catalogue for conifer genome analyses. Plant Physiology 157: 14–28. 10.1104/pp.111.179663

55. Rimmer A, Phan H, Mathieson I, Iqbal Z, Twigg SRF, WGS500 Consortium, Wilkie AOM, McVean G, Lunter G. 2014. Integrating mapping-, assembly-and haplotype-based approaches for calling variants in clinical sequencing applications. Nature Genetics 46: 912–918. 10.1038/ng.3036

56. Salse J. 2012. In silico archeogenomics unveils modern plant genome organisation, regulation and evolution. Curr. Opin. Plant Biol. 15: 122–1. 10.1016/j.pbi.2012.01.001

57. Selvaraj S, R Dixon J, Bansal V, Bing R. 2013. Whole-genome haplotype reconstruction using proximity-ligation and shotgun sequencing. Nat Biotechnol 31:1111–1118. 10.1038/nbt.2728

58. Sewell MM, Sherman BK, Neale DB. 1999. A consensus map for loblolly pine (Pinus taeda L.). I. Construction and integration of individual linkage maps from two outbred three-generation pedigrees. Genetics 151(1):321–30. 10.1093/genetics/151.1.321

59. Song Q, Jenkins J, Jia G, Hyten DL, Pantalone V, Jackson SA, Schmutz J, Cregan PB. 2016. Construction of high-resolution genetic linkage maps to improve the soybean genome sequence assembly Glyma1.01. BMC Genom. 17:33. 10.1186/s12864-015-2344-0

60. Stival Sena J, Giguère I, Rigault P, Bousquet J, Mackay J. 2018. Expansion of the dehydrin gene family in conifers is associated with considerable structural diversity and drought responsive expression. Tree Physiology 38 (3):442–456. 10.1093/treephys/tpx125

61. Stival Sena, J, Giguère I, Boyle B, Rigault P, Birol I, Zuccolo A, Ritland K, Ritland C, Bohlmann J, Jones S, Bousquet J, MacKay J. 2014. Evolution of gene structure in the conifer Picea glauca: a comparative analysis of the impact of intron size. BMC Plant Biology 14:95. 10.1186/1471-2229-14-95

62. Sturtevant AH. 1913. A Third Group of Linked Genes in Drosophila Ampelophila. Science 37:990– 992. 10.1126/science.37.965.9

63. Tumas HR, Soufi Z, Woolliams JA, McLean JP, Lee S, Cottrell JE, Ilska JJ, Lopez G, MacKay J. 2021. Stranger in a strange land: genetic variation of native insect resistance biomarkers in UK Sitka spruce (Picea sitchensis [Bong.] Carr.). Forestry: An International Journal of Forest Research 94(5):734–744. 10.1093

64. Tuskan GA, Difazio S, Jansson S, Bohlmann J, Grigoriev I, Hellsten U, Putnam N, Ralph S, Rombauts S, Salamov A, Schein J, Sterck L, Aerts A, Bhalerao RR, Bhalerao RP, Blaudez D, Boerjan W, Brun A, Brunner A, Busov V, Campbell M, Carlson J, Chalot M, Chapman J, Chen GL, Cooper D, Coutinho PM, Couturier J, Covert S, Cronk Q, Cunningham R, Davis J, Degroeve S, Déjardin A, Depamphilis C, Detter J, Dirks B, Dubchak I, Duplessis S, Ehlting J, Ellis B, Gendler K, Goodstein D, Gribskov M, Grimwood J, Groover A, Gunter L, Hamberger B, Heinze B, Helariutta Y, Henrissat B, Holligan D, Holt R, Huang W, Islam-Faridi N, Jones S, Jones-Rhoades M, Jorgensen R, Joshi C, Kangasjärvi J, Karlsson J, Kelleher C, Kirkpatrick R, Kirst M, Kohler A, Kalluri U, Larimer F, Leebens-Mack J, Leplé JC, Locascio P, Lou Y, Lucas S, Martin F, Montanini B, Napoli C, Nelson DR, Nelson C, Nieminen K, Nilsson O, Pereda V, Peter G, Philippe R, Pilate G, Poliakov A, Razumovskaya J, Richardson P, Rinaldi C, Ritland K, Rouzé P, Ryaboy D, Schmutz J, Schrader J, Segerman B, Shin H, Siddiqui A, Sterky F, Terry A, Tsai CJ, Uberbacher E, Unneberg P, Vahala J, Wall K, Wessler S, Yang G, Yin T, Douglas C, Marra M, Sandberg G, Van de Peer Y, Rokhsar D. 2006. The genome of black cottonwood, Populus trichocarpa (Torr. & Gray). Science 313(5793):1596–604. 10.1126/science.1128691

65. Van Ghelder C, Parent GJ, Rigault P, Prunier J, Giguère I, Caron S, Stival Sena J, Deslauriers A, Bousquet J, Esmenjaud D, MacKay J. 2019. The large repertoire of conifer NLR resistance genes includes drought responsive and highly diversified RNLs. Scientific Reports 9: 1–13. 10.1038/s41598-019-47950-7

66. Velmurugan J, Mollison E, Barth S, Marshall D, Milne L, Creevey CJ, Lynch B, Meally H, McCabe M, Milbourne, D. 2016. An ultra-high-density genetic linkage map of perennial ryegrass (Lolium perenne) using genotyping by sequencing (GBS) based on a reference shotgun genome assembly. Ann. Bot. 118:71–87. 1010.1093/aob/mcw081

67. Warren RL, Keeking CI, Yuen MMS, Raymond A, Taylor GA, Vandervalk BP, Hamid Mohamadi H, Paulino D, Chiu R, Jackman SD, Robertson G, Yang C, Boyle B, Hoffmann M, Weigel D, Nelson DR, Ritland C, Isabel N, Jaquish B, Yanchuk A, Bousquet J, Jones SJM, MacKay J, Birol I, Bohlmann J. 2015. Improved white spruce (Picea glauca) genome assemblies and annotation of large gene families of conifer defense metabolism. The Plant Journal 83:189–212. 10.1111/tpj.12886

68. Westbrook JW, Chhatre VE, Wu LS, Chamala S, Neves LG, Muñoz P, Martínez-García PJ, Neale DB, Kirst M, Mockaitis K, Nelson CD, Peter GF, Davis JM, Echt CS. 2015. A Consensus Genetic Map for Pinus taeda and Pinus elliottii and Extent of Linkage Disequilibrium in Two Genotype-Phenotype Discovery Populations of Pinus taeda. G3 (Bethesda) 5(8):1685-94. 10.1534/g3.115.019588

69. Wickham H. 2016.. ggplot2: Elegant Graphics for Data Analysis. Springer-Verlag New York. ISBN 978-3-319-24277-4, https://ggplot2.tidyverse.org.

70. Xiong X, Gou J, Liao Q, Li Y, Zhou Q, Bi G, Li C, Du R, Wang X, Sun T, Guo L, Liang H, Lu P, Wu Y, Zhang Z, Ro D-K, Shang Y, Huang S, Yan J. 2021. The Taxus genome provides insights into paclitaxel biosynthesis. Nat. Plants 7:1026–1036. 10.1038/s41477-021-00963-5

71. Xu X, Pan S, Cheng S, Zhang B, Mu D, Ni P, Zhang G, Yang S, Li R, Wang J, Orjeda G, Guzman F, Torres M, Lozano R, Ponce O, Martinez D, De la Cruz G, Chakrabarti SK, Patil VU, Skryabin KG, Kuznetsov BB, Ravin NV, Kolganova TV, Beletsky AV, Mardanov AV, Di Genova A, Bolser DM, Martin DM, Li G, Yang Y, Kuang H, Hu Q, Xiong X, Bishop GJ, Sagredo B, Mejía N, Zagorski W, Gromadka R, Gawor J, Szczesny P, Huang S, Zhang Z, Liang C, He J, Li Y, He Y, Xu J, Zhang Y, Xie B, Du Y, Qu D, Bonierbale M, Ghislain M, Herrera Mdel R, Giuliano G, Pietrella M, Perrotta G, Facella P, O’Brien K, Feingold SE, Barreiro LE, Massa GA, Diambra L, Whitty BR, Vaillancourt B, Lin H, Massa AN, Geoffroy M, Lundback S, DellaPenna D, Buell CR, Sharma SK, Marshall DF, Waugh R, Bryan GJ, Destefanis M, Nagy I, Milbourne D, Thomson SJ, Fiers M, Jacobs JM, Nielsen KL, Sønderkær M, Iovene M, Torres GA, Jiang J, Veilleux RE, Bachem CW, de Boer J, Borm T, Kloosterman B, van Eck H, Datema E, Hekkert Bt, Goverse A, van Ham RC, Visser RG. 2011. Genome sequence and analysis of the tuber crop potato. Nature 475:189–195. 10.1038/nature10158

72. Zimin A, Stevens KA, Crepeau MW, Holtz-Morris A, Koriabine M, Marçais G, Puiu D, Roberts M, Wegrzyn JL, de Jong PJ, Neale DB, Salzberg SL, Yorke JA, Langley CH. 2014. Sequencing and Assembly of the 22-Gb Loblolly Pine Genome. Genetics 196(3):875–890. 10.1534/genetics.113.159715

